# Estimating the frequency of virulence against an *Stb* gene in *Zymoseptoria tritici* populations by bulk phenotyping on checkerboard microcanopies of wheat NILs

**DOI:** 10.1101/2023.12.18.572116

**Authors:** Frédéric Suffert, Stéphanie Le Prieur, Sandrine Gélisse, Emmie Dzialo, Cyrille Saintenac, Thierry C. Marcel

## Abstract

Monitoring virulent strains within fungal pathogen populations is crucial to improve host resistance deployment strategies. Such monitoring increasingly involves field pathogenomics studies of molecular polymorphisms in genomes based on high-throughput screening technologies. However, it is not always straightforward to predict virulence phenotypes from these polymorphisms and *in planta* phenotyping remains necessary. We developed a method for ‘bulk phenotyping on checkerboard microcanopies of wheat near-isogenic lines’ (BPC) for estimating the frequency of virulence against an *Stb* gene in populations of *Zymoseptoria tritici*, the causal agent of *Septoria tritici* blotch in wheat, without the need for strain-by-strain phenotyping. Our method involves the uniform inoculation of a microcanopy of two wheat lines – one with the resistance gene and the other without it – with a multi-strain cocktail representative of the population to be characterized, followed by the differential quantification of infection points (lesions). Using *Stb16q*, a resistance gene that has recently broken down in Europe, we found a robust correlation between the ratio of the mean number of lesions on each wheat line and the frequency of virulent strains in the inoculum. Using pairs of virulent and avirulent strains, and synthetic populations consisting of 10 virulent strains and 10 avirulent strains mixed in different proportions, we validated the principle of the method and established standard curves at virulence frequencies close to those observed in natural conditions. We discuss the potential of this method for virulence monitoring in combination with recently developed molecular methods.

## Introduction

The fungus *Zymoseptoria tritici* is the causal agent of *Septoria tritici* blotch (STB), a disease responsible for major yield losses on bread wheat in north-western Europe (Fones & Gurr, 2015). Field populations of *Z. tritici* are extremely large and genetically diverse, due to recurring annual cycles of sexual reproduction (McDonald et al., 2022; Suffert et al., 2011). These characteristics account for the ability of the pathogen to adapt to biotic and abiotic environmental conditions and to counter protective strategies based on resistant varieties and fungicides. Comparisons of neutral allele frequencies and the use of ‘low-density’ molecular markers have shown high degrees of similarity between populations at spatial scales ranging from the field to continents (Linde et al., 2002; Zhan et al., 2003), whereas ‘high-density’ genotyping (complete genome resequencing) has recently revealed a genetic structure tracing the evolutionary trajectory of *Z. tritici* populations at a global scale (Feurtey et al., 2023). Moreover, despite the apparent lack of differentiation of populations over very large spatial scales, *Z. tritici* can adapt to selective pressures exerted locally (Suffert et al., 2015). The effective management of STB is dependent on identification of determinants of these adaptations and the development of phenotypic and genotypic methods for monitoring them in the pathogen populations.

One of the most sustainable management strategies for controlling STB is the use of resistant varieties. Resistance is governed by quantitative trait loci (QTLs) and major genes (*Stb*). The pathogenicity of *Z. tritici* mirrors host resistance, with qualitative and quantitative components (Brading et al., 2002; Gohari, 2015; Stewart et al., 2016), both of which may participate in gene-for-gene interactions with wheat resistance genes (Meile et al., 2017; Zhong et al., 2017; Brading et al., 2002; Langlands-Perry et al., 2023). In such interactions, avirulence effector proteins from the fungus interact with plant disease resistance proteins to trigger a resistance response (Flor, 1956). To date, 23 *Stb* genes have been genetically mapped in wheat (Brown et al., 2015; Yang *et al*., 2018; Langlands-Perry et al., 2022), but none of these genes is effective against all strains of *Z. tritici*. Some *Stb* genes have been present in commercial varieties over long periods (e.g. *Stb6*), whereas others were introduced only recently, into a small number of varieties (e.g. *Stb16q*; Brown et al., 2015). Breeding programs improve the general level of resistance of newly released varieties, as illustrated by the increase in the mean resistance level of varieties deployed in France from 2005 to 2020 (du Cheyron, 2020). Nevertheless, in certain wheat areas, the low genetic diversity of the crops grown exerts a unidirectional selection pressure on *Z. tritici* populations, fostering the breakdown of the resistance genes carried by these varieties and resulting in a rapid increase in the frequency of the virulent strains able to overcome these resistances (Brown & Tellier, 2011). The occurrence of virulence within local pathogen populations varies significantly between resistance genes and between geographic regions. The frequency of virulence is influenced by several factors, including the prevalence of cultivated wheat varieties carrying a specific resistance gene and the commercial lifecycle of these varieties, encompassing the duration of their deployment and the extent to which they are adopted.

Virulence monitoring at different spatiotemporal scales is crucial for the reasoned deployment of resistance genes to prevent their rapid breakdown. The development and use of suitable molecular markers is a powerful approach, but may not necessarily be suitable for monitoring the emergence of a new virulence, which is generally detected in the field when the first symptoms appear on a resistant variety (i.e. when the resistance breakdown is already underway). This was the case for the virulence on the *Stb16q* gene in France (Orellana-Torrejon, 2022a) and Ireland (Kildea et al., 2020) in the late 2010s. In practice, virulence can be monitored at an earlier stage, by sampling a few infected leaves from surveyed fields and varieties, obtaining monospore isolates and performing individual (strain-by-strain) *in planta* phenotyping on a set of differential wheat lines (i.e. carrying different resistance genes). However, detecting changes in virulence frequencies by individual phenotyping over a long period of time or in analytical annual field experiments is challenging, due to the large number of strains to be tested (e.g. Orellana-Torrejon et al., 2022a, 2022b). It would be quicker to phenotype a group of strains simultaneously to determine the frequency of virulent individuals, but this approach is difficult to implement, due to the inoculation method and the characteristics of disease expression. It is tempting to hypothesize that the ratio of symptom expression between plants with and without a resistance gene is correlated with frequency of virulent strains in a bulk. If sufficiently robust, such a correlation could be used to estimate virulence frequencies at population level. The principle behind this approach is that all strains can infect a ‘susceptible’ plant, whereas only virulent strains can infect a plant carrying the corresponding resistance gene. Counting the number of points of infection on each type of plant and calculating the difference between plants with and without the resistance gene would therefore make it possible to estimate the proportion of virulent strains. This would require the inoculation of each plant with an identical pathogen population (i.e. the bulk) – ideally in a simultaneous inoculation event – with well-delimited, or only slightly coalescing lesions on each plant type. To our knowledge, this principle has never before been applied in this way in plant pathology under the conditions mentioned above. Groth and Ozmon (2002) implemented a slightly different approach for a rust fungus and methods based on this principle have been suggested for foodborne pathogens (Harrand et al., 2019). We describe here the development of a method based on this principle, using STB as a case study.

## Materials and methods

### Plant material

All experiments were performed with two near-isogenic lines (NILs) of bread wheat (*Triticum aestivum*) carrying either no *Stb* gene (NIL^stb^) or *Stb16q* (NIL^Stb16q^). These two NILs were developed as described by Battache et al. (2021), from Chinese Spring (CS) carrying *Stb6* (Chartrain et al., 2005) and the synthetic hexaploid wheat line TA4152-19 carrying *Stb16q* (Saintenac et al., 2021). Four seeds of the same line – NIL^stb^ or NIL^Stb16q^ – were sown in 0.4-liter pots (7 × 7 × 8 cm) to allow for possible emergence failure and to ensure that we obtained three seedlings per pot. During the course of all experiments, the pots were kept in a growth chamber under a controlled light/dark cycle (16 hours of light at 300 µmol.m^-2^.s^-1^ – Osram Lumilux L58W/830 – at 22°C and 80% RH / 8 hours of dark at 18°C and 100% RH). Fifteen days after sowing, a 5-cm long section was drawn with a black marker pen in the middle of the second leaf of each seedling. Eight pots of each line were combined in a staggered pattern in a dish to form a regularly heterogeneous 30-cm-square checkerboard microcanopy (Figure 1).

**Figure 1.**
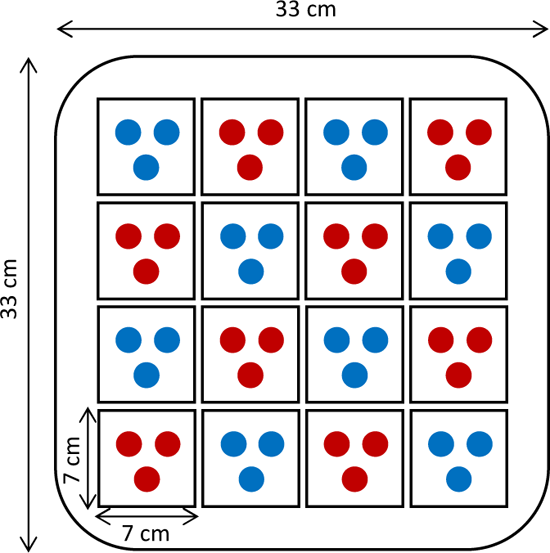
Experimental design showing the checkerboard microcanopy. Each dot represents a wheat seedling (blue for NIL^stb^, red for NIL^Stb16q^).

### Fungal material

We used different *Z. tritici* strains collected from cv Apache (moderately susceptible to STB) and Cellule (carrying *Stb16q*) in a field trial in Thiverval-Grignon (France) in 2018 and 2019 (Table 1). The virulence status of these strains against *Stb16q* (qualitative component of pathogenicity, here assessed as the ability to cause symptoms on cv Cellule seedlings; ‘vir’ for virulent and ‘avr’ for avirulent) and their aggressiveness (quantitative component of pathogenicity on a susceptible host, here expressed as the percentage of the leaf area displaying sporulation on Apache seedlings) had already been determined (Orellana-Torrejon et al., 2022a). Different mixtures of vir and avr strains were used in three complementary experiments to evaluate and validate the ‘bulk phenotyping on a checkerboard microcanopy’ method (hereafter called ‘BPC’): two binary vir-avr mixtures (Mix 1A, Mix 1B), a synthetic population consisting of 10 vir and 10 avr strains (Mix 2), and an even more diverse population resulting from a sexual cross between a vir and an avr strain (Mix 3) (Table 1).

**Table 1.**
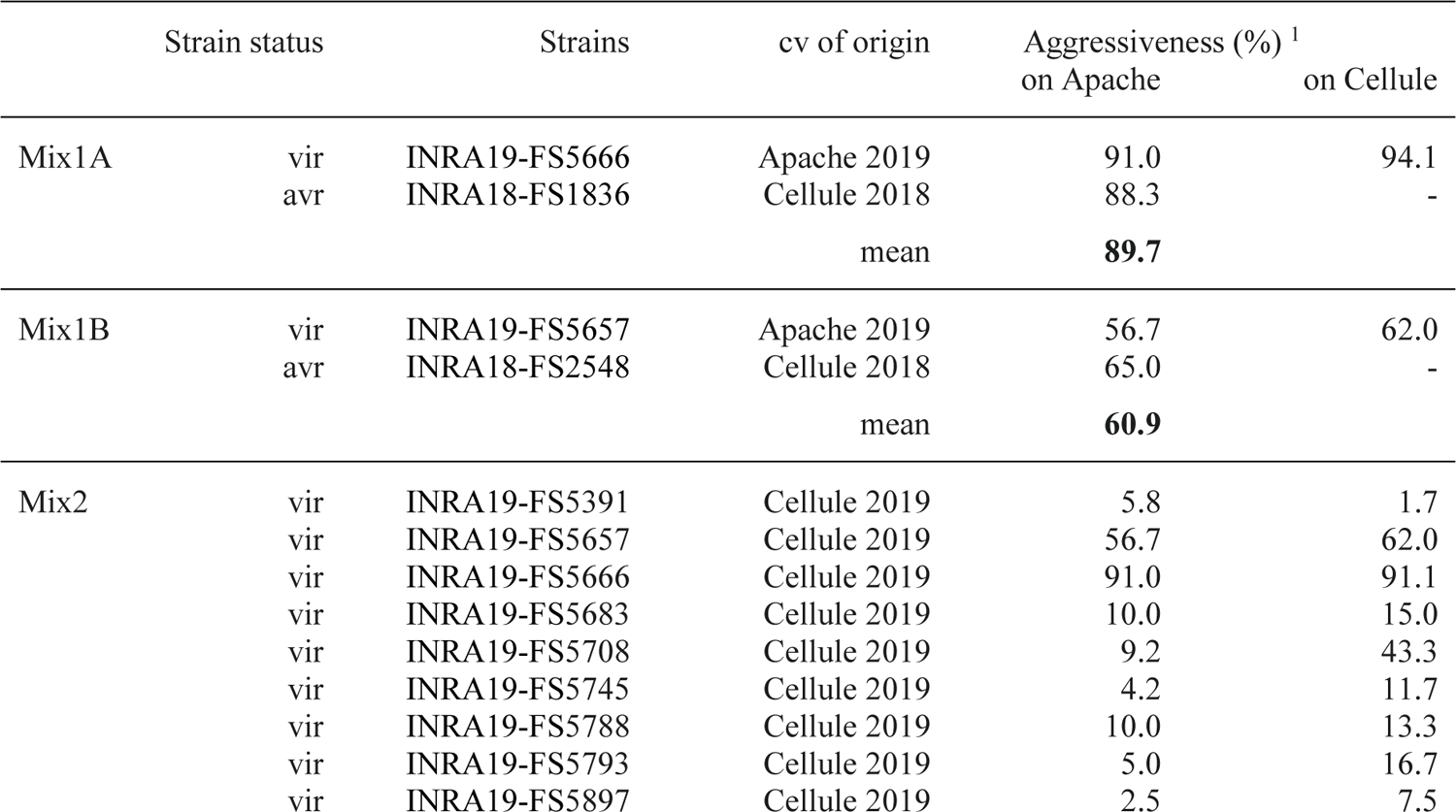
*Zymoseptoria tritici* strains used in different mixtures (Mix1A, Mix1B, Mix2 and Mix3) for development and validation of the BPC method.

### Experimental design

A first experiment was performed to test the hypotheses and expectations on which the BPC method was conceived, to assess its practicality and to estimate its sensitivity to inoculum concentration. We established 11 checkerboard microcanopies, which we inoculated with a suspension containing a mixture of one virulent (vir) and one avirulent (avr) *Z. tritici* strain in various proportions, from 0 to 1, in 0.10 increments. Preliminary tests showed that inoculum concentration was crucial for the accuracy and efficacy of the method (Figure 2). A concentration of 10^4^ spores.mL^-1^ generated very few sporadic lesions, whereas a concentration of 10^6^ spores.mL^-1^ generated very large areas of sporulation, making it difficult to count lesions due to their coalescence. A concentration of 10^5^ spores.mL^-1^ therefore appeared to be the most appropriate order of magnitude for these experiments. We therefore used 1.0 × 10^5^, 1.5 × 10^5^ and 2.0 × 10^5^ spores.mL^-1^ in the first experiment. This experiment was repeated once, to obtain two replicates with two pairs of vir and avr strains selected on the basis of similar levels of aggressiveness within a pair: high aggressiveness (89.7% on average) for Mix1A (FS5666 + FS1836) and moderate aggressiveness (60.9% on average) for Mix1B (FS5657 + FS2548) (Table 1).

**Figure 2.**
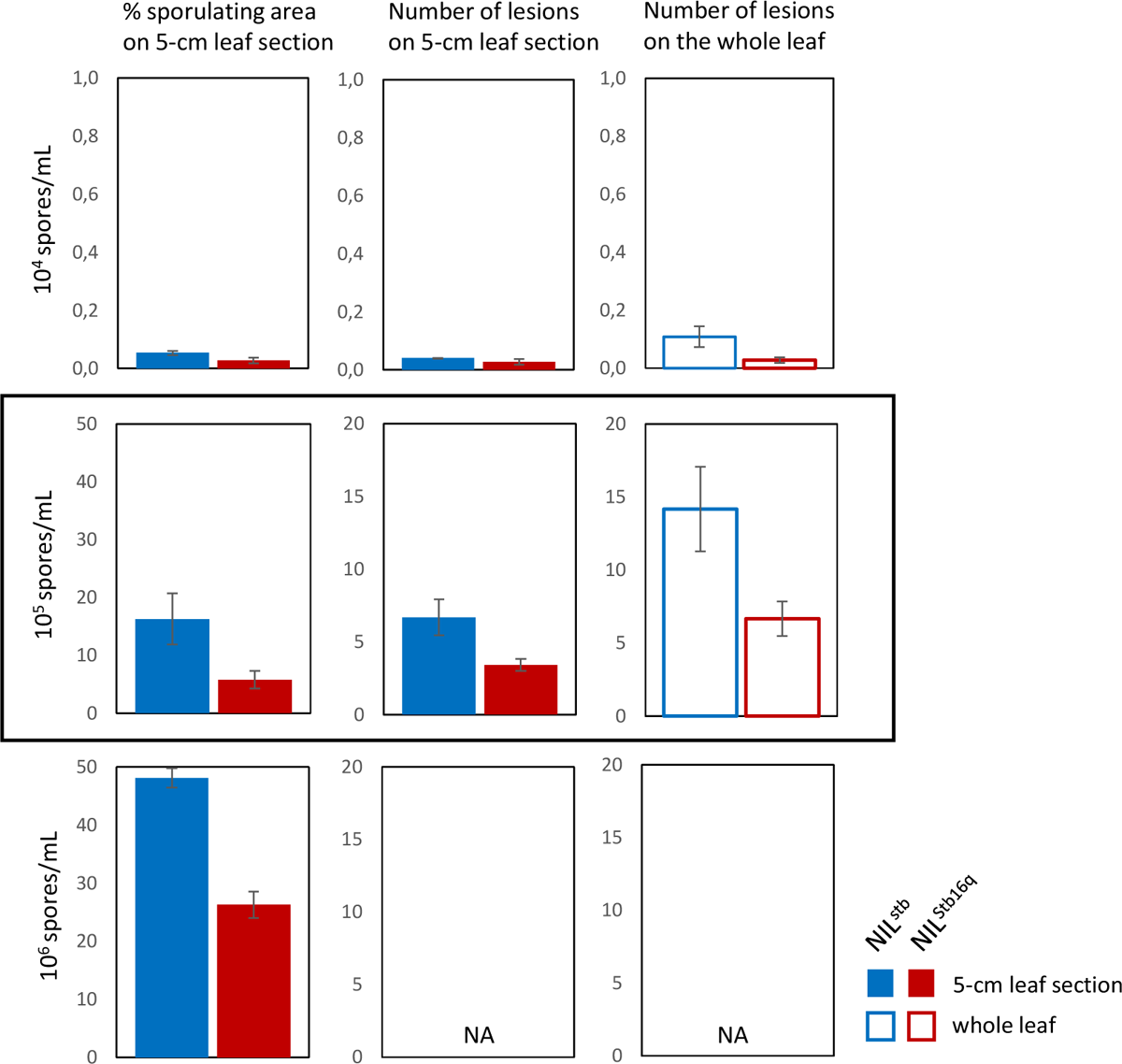
Results of preliminary tests to determine the most relevant *Zymoseptoria tritici* inoculum concentration (10^4^ spores.mL^-1^, 10^5^ spores.mL^-1^ or 10^6^ spores.mL^-1^) for the development of the BPC method with the two near-isogenic lines, NIL^stb^ (blue) and NIL^Stb16q^ (red), of bread wheat. Disease assessments were performed 18 days post inoculation (dpi, data shown here) and 21 dpi (data not shown). NA indicates that it was not possible to count the individual lesions for a concentration of 10^6^ spores.mL^-1^ because the high inoculum pressure induced rapid and complete leaf necrosis with the coalescence of most lesions.

A second experiment was conducted to validate the BPC method across a spectrum of proportions of the low-virulence strain (0, 0.05, 0.10, 0.20, and 0.50), with a more diverse pathogen population consisting of 10 vir and 10 avr *Z. tritici* strains (Mix2) in suspension at an overall concentration of 2.0 × 10^5^ spores.mL^-1^. The 20 strains were chosen to encompass a spectrum of aggressiveness (Table 1) and to mirror the diverse composition of pathogen populations observed in natural field conditions, although the genotypic diversity of *Z. tritici* is known to be much greater (McDonald et al., 2022). This second experiment involved three checkerboard microcanopies for each proportion, with one repetition, to obtain two replicates, to estimate the variability of the response.

A third experiment was conducted to apply the BPC method by characterizing an even more diverse population consisting of a large number of *Z. tritici* progenies from a sexual cross between a vir and an avr strain (Table 1). The precise composition of the tested population was unknown, but the vir/avr ratio was expected to tend to 1 as the genetic determinism of virulence against *Stb16q* is monogenic and involves a gene-for-gene interaction (Saintenac et al., 2021). The progeny was obtained as described by Suffert et al. (2016) from cv Apache wheat plants co-inoculated with the strains FS3098 (vir) and FS2851 (avr). Colonies derived from ascospores were obtained in Petri dishes containing PDA following the ejection of ascospores from wheat residues (see Supplementary Figure in Orellana-Torrejon et al., 2023c). These colonies, originating from more than 30 clusters representing a diverse array of a few hundred different *Z. tritici* genotypes, were scraped up and mixed immediately after collection (Mix3). This application of the BPC method, through the inoculation of one microcanopy with Mix3 and another with Mix1 as a control, was repeated once, to obtain two replicates.

### Inoculation procedure

All the inocula used in the experiment were obtained from blastospores that had been stored at −80°C. Each strain was cultured in a Petri dish containing PDA (potato dextrose agar, 39 g.L^-1^) placed at 18°C in the dark. This operation was repeated one week before inoculation, to make it possible to work with homogeneous subcultures. Blastospore suspensions were prepared by flooding the surface of the six-day-old subcultures with sterile distilled water and then scraping the agar surface with a sterilized glass rod as described by Suffert et al. (2013). Single-isolate stock solutions were then prepared by adjusting the blastospore suspension to the required concentration with a Malassez cell.

For the first experiment, each vir or avr strain suspension (Table 1) was adjusted to a concentration of 1.0 × 10^5^, 1.5 × 10^5^, and 2.0 × 10^5^ spores.mL^-1^ and the vir and avr suspensions were then mixed together in the appropriate volumes to obtain the required proportions of each strain. Each inoculum suspension used in the second experiment was obtained by mixing the appropriate volumes of a suspension of 10 avr strains in equal proportions (obtained by mixing 20 mL of each of the 10 corresponding suspensions adjusted individually to a concentration of 2.0 × 10^5^ spores.mL^-1^) with a similar suspension consisting of 10 vir strains. The inoculum suspension used in the third experiment was prepared by subculturing the multistrain stock of blastospores and adjusting the concentration to 2.0 × 10^5^ spores.mL^-1^. Suspensions were prepared the day before inoculation and stored at 4°C after the addition of two drops of Tween 20 (0.1 % v/v; Sigma).

All inoculum suspensions were applied with an atomizer (Ecospray, VWR, France) 16 days after sowing. Each checkerboard microcanopy was turned and sprayed with criss-cross movements for 45 s to ensure a homogeneous distribution of microdroplets on all the leaves (Figure 3A). The amount of suspension sprayed (20 ± 1 mL) was determined by weighing each flask at the start and end of the application. Each checkerboard microcanopy was bagged for three days immediately after spraying, to promote infection (Figure 3B). It was then kept in the growth chamber until disease assessment. The second half of the third and fourth leaves, which had not yet emerged or had only partially emerged at the time of inoculation, were cut off 12 days after inoculation (dpi) to clear the microcanopies. Plants were watered as needed (three times per week) for the entire duration of the experiment.

**Figure 3.**
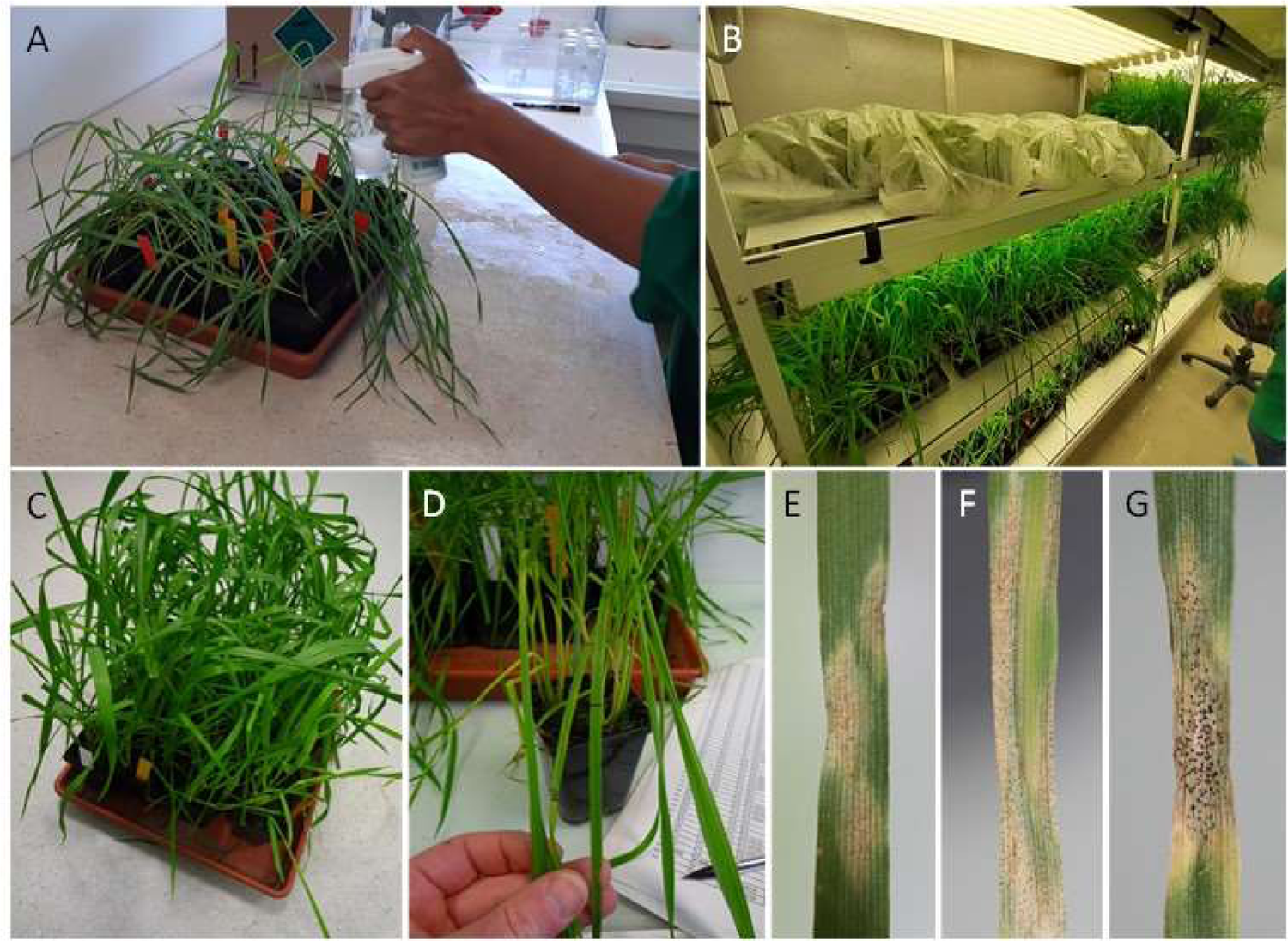
Experimental steps in the BPC method. A. Application, with an atomizer, of a *Z. tritici* blastospore suspension on a microcanopy 16 days after sowing. B. Bagged checkerboard microcanopy. C. A checkerboard microcanopy 21 days after inoculation. D. Disease assessment on both the 5-cm section and the whole second leaf of each seedling. E-F-G. Three leaves with STB lesions, from the least to the most coalescent, representative of the symptoms observed.

### Disease assessment

STB lesions were counted to estimate the number of infection points on both the 5-cm section and the whole area of the second leaf of each inoculated seedling at 21 dpi in the first experiment and 19 dpi in the second and third experiments (Figure 3C-G). Preliminary tests had shown that earlier disease assessment missed a number of lesions that emerged late, whereas later assessment was problematic because of the coalescence of some lesions due to the desiccation of the most severely attacked leaves.

In some sporulating areas occupied by coalescent lesions (especially in the terminal part of the leaves, which were thin and often curled in cases of severe attack), the number of infection points was estimated based on the mean area of a lesion. We found that it became difficult to count individual lesions if there were more than 20-30 lesions on the 5-cm section (the widest part of the leaf) and 50-60 lesions on the whole leaf. These maximum counting values were assigned to the most severely attacked leaves (sporulating surface area tending to 100%). This implies that a lesion occupies about 2% of the whole leaf area and that the 5-cm section represented 40% of the total leaf area, at the time of inoculation at least. We are aware that this is a rough estimate, given the differences in surface area between leaves, but it makes it possible to compare the effects of different treatments and different variables. Differences were observed in the timing of lesion appearance, with some lesions appearing younger than others. We therefore felt it was more relevant to count the lesions than to estimate a sporulating area as we usually do in studies comparing aggressiveness between strains (e.g. Orellana-Torrejon et al., 2022a) or assessing the effect of environmental factors on disease development (e.g. Boixel et al., 2022).

## Results

### Respect of basic assumptions

The first experiment was used to assess the validity of the principle of the BPC method by checking that the basic assumptions were satisfied and validating the most relevant experimental conditions. An ANOVA performed on the data from this first experiment (see Supplementary_material_full_dataset) showed that the effects of wheat line, concentration of inoculum suspension and vir/avr proportion in the inoculum suspension were significant, unlike that of mixture (Mix1A, Mix1B) (Table S1).

A first important outcome was that inoculum suspensions containing only an avr strain (FS1836 or FS2548 in the first experiment) caused no symptoms on NIL^Stb16q^ (Figure 4). This was confirmed in the second experiment after inoculation with the mixture containing only the 10 avr strains (see below; Figure 8A). These findings demonstrate that all the avr strains used here were, indeed, avirulent in the experimental conditions used and that the procedures followed, including inoculation and incubation in the growth chamber in particular, did not lead to contamination with the vir strains present in the other inoculum suspensions.

**Figure 4.**
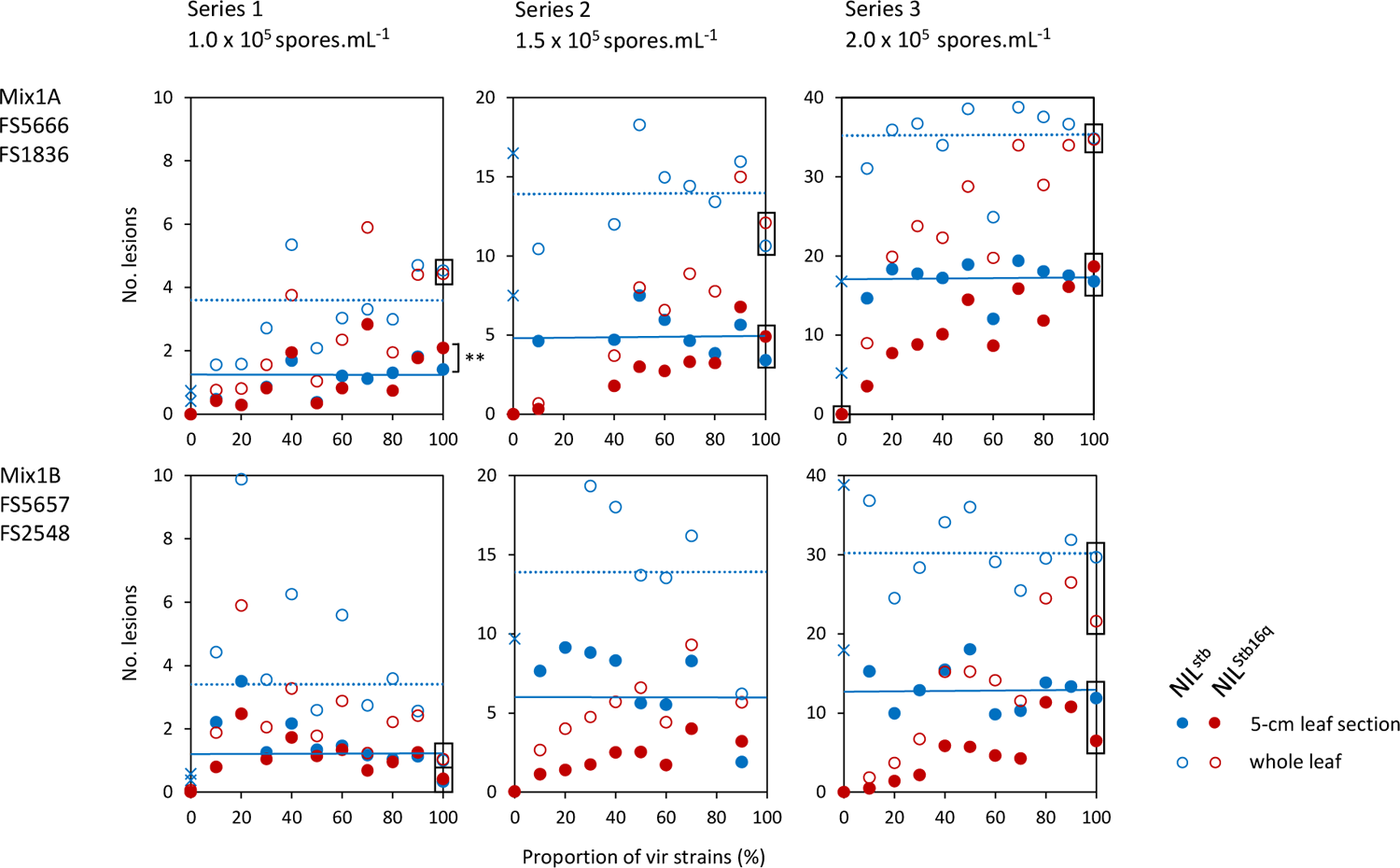
Mean number of STB lesions counted on the 5-cm leaf section and on the whole leaf of wheat lines NIL^stb^ (blue) and NIL^Stb16q^ (red) in a microcanopy according to the proportion of vir strains in the inoculum suspension (vir FS5666 and avr FS1836 for Mix1A; vir FS5657 and avr FS2548 for Mix1B) and the concentration of the inoculum suspension (1.0 × 10^5^ spores.mL^-1^, 1.5 × 10^5^ spores.mL^-1^ or 2.0 × 10^5^ spores.mL^-1^). Each point is the mean number of lesions on 24 marked leaves. Horizontal blue lines represent the mean number of lesions appearing on NIL^stb^ regardless of the proportion of vir strains in the inoculum suspension.

A second major finding was the similarity of the quantitative response obtained in analogous, compatible interactions: (i) the avr and vir strains induced disease severity of the same order of magnitude (number of lesions) on NIL^stb^, as highlighted by the moderate differences in the number of lesions on NIL^stb^ between the various vir/avr proportions tested (horizontal blue lines in Figure 4) especially for Mix1A Series3 (overall non-significant effect of vir/avr proportion on the mean number of STB lesions counted on the 5-cm leaf section; Kruskal-Wallis test; Suppl. Table S2) relative to the differences observed for the various proportions on NIL^Stb16q^ (linear model of the mean number of STB lesions counted on the 5-cm leaf section against vir/avr proportion; Suppl. Table S3); (ii) there was no significant difference in the numbers of lesions obtained between NIL^stb^ and NIL^Stb16q^ inoculated with only vir strains (see vertical boxes in Figure 4), except for Series1 of Mix1A when lesions were counted on the 5-cm leaf section. This second outcome is consistent with the similar levels of aggressiveness of the strains in Mix1A (around 90%) and of those in Mix1B (around 60%) estimated by Orellana-Torrejon et al. (2022a).

### Effect of inoculum concentration

In the first experiment we confirmed that the concentration of the inoculum suspension (here 10^5^ spores.mL^-1^) has a significant impact on disease severity and must therefore be chosen carefully (see preliminary tests presented in Figure 2). The number of lesions increased proportionally with the concentration of the inoculum suspension: the mean number of lesions on NIL^stb^ for Mix1A and Mix1B was 1.31 on a 5-cm leaf section for 1.0 × 10^5^ spores.mL^-1^, 5.97 for 1.5 × 10^5^ spores.mL^-1^ and 15.07 for 2.0 × 10^5^ spores.mL^-1^ (Figure 4).

Moreover, disease scores for 5-cm leaf sections and whole leaves were strongly correlated (degree 3 polynomial, r^2^ = 0.98; Figure 5), regardless of the wheat line (NIL^stb^ or NIL^Stb16^). The relationship between the two scores was proportional for low to moderate lesion densities, corresponding to a mean of fewer than 12 lesions on a 5-cm leaf section (equivalent of 25 lesions on a whole leaf). At very high disease severities, with lesion densities above this threshold, visual estimates of the number of lesions became less reliable because of a saturation/coalescence effect. The relationship was no longer fully proportional, as shown by the flattening of the curve. Disease assessment on a 5-cm leaf section makes it possible to standardize the results, as leaf size is variable. It also makes the disease assessment quicker, but it requires the prior marking of the leaves.

**Figure 5.**
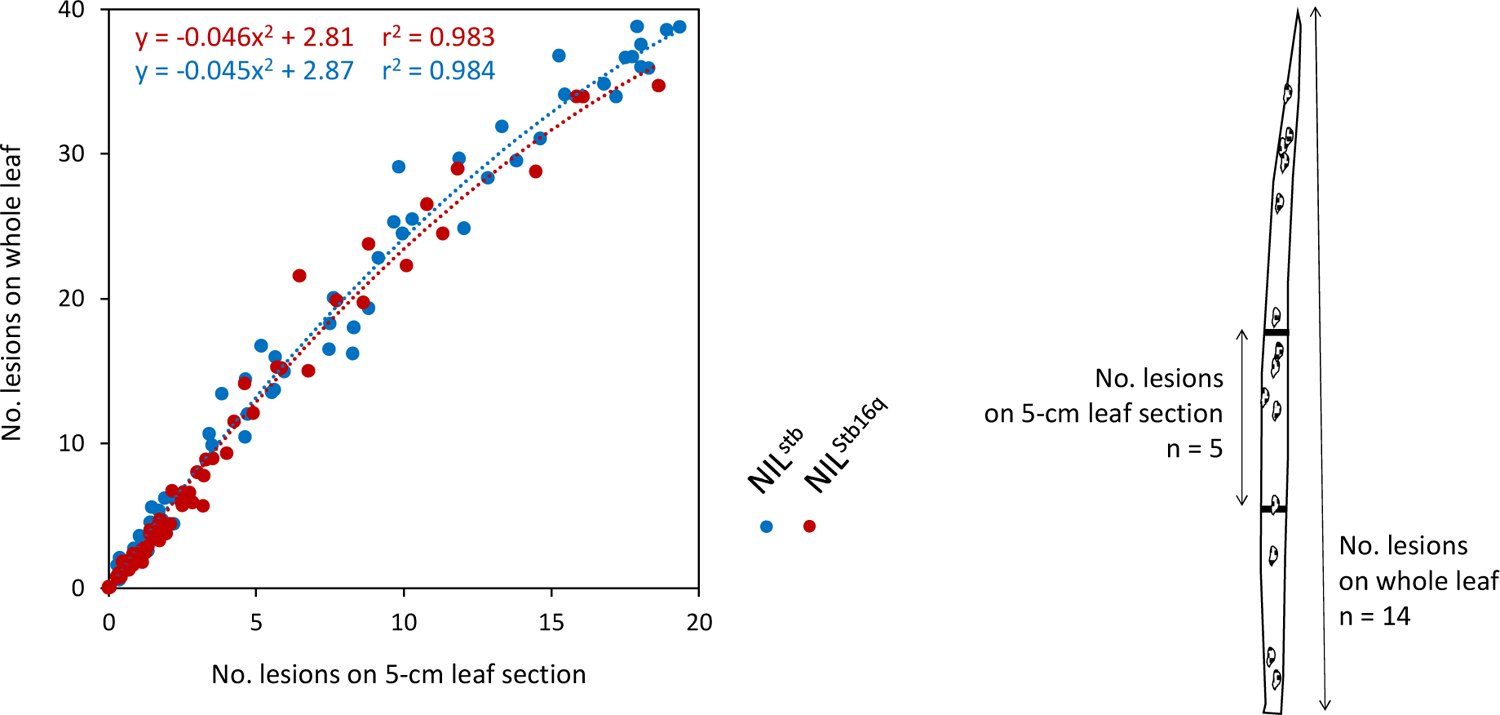
Correlation between the number of STB lesions counted on the 5-cm leaf section and on the whole leaf for a given wheat line (NIL^stb^ in blue; NIL^Stb16^ in red) in a microcanopy. Each point is the mean number of lesions on 24 marked leaves (pooled data from Mix1A and Mix1B in the first experiment and for Mix2 in the second experiment).

The ratio of the mean number of STB lesions counted on line NIL^Stb16q^ to that on NIL^stb^ (ρ_L_) increased from 0 to 1 with an increased proportion of vir strains in the inoculum suspension (Figure 6). The correlation between ρ_L_ and the proportion of vir strains was poor at an inoculation concentration of 1.0 × 10^5^ spores.mL^-1^ (r^2^ = 0.194 for Mix 1A and 0.681 for Mix 1B, with assessments performed on the 5-cm leaf section), but strong at inoculum concentrations of 1.5 × 10^5^ spores.mL^-1^ (r^2^ = 0.907 and 0.664) and 2.0 × 10^5^ spores.mL^-1^ (r^2^ = 0.876 and 0.827), suggesting that virulence frequency can be estimated by the BPC method with these last two concentrations.

**Figure 6.**
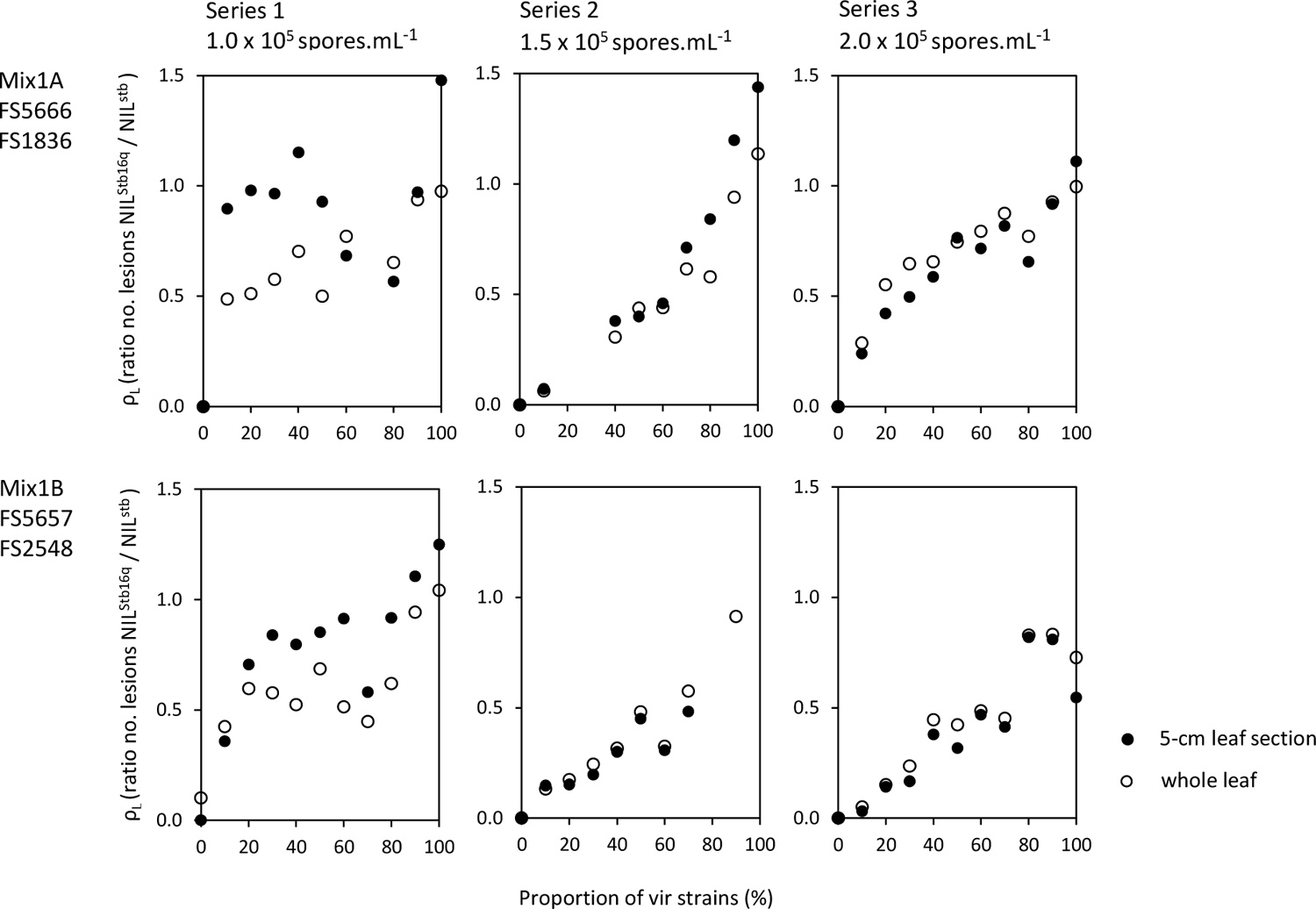
Ratios (ρ_L_) of the mean number of STB lesions counted on the 5-cm leaf section (black points) or on the whole leaf (white points) of wheat line NIL^Stb16q^ relative to and NIL^stb^ in a microcanopy according to the proportion of vir strains in the inoculum suspension (vir FS5666 and avr FS1836 for Mix1A; vir FS5657 and avr FS2548 for Mix1B) and the concentration of the inoculum suspension (1.0 × 10^5^ spores.mL^-1^, 1.5 × 10^5^ spores.mL^-1^ and 2.0 × 10^5^ spores.mL^-1^).

### Establishment of standard curves

‘Standard curves’ linking ρ_L_ and the proportion of vir strains in the mixture, intended to be used for estimating the proportion of virulence within a pathogen population, were established by pooling the data set obtained with inoculum concentrations of 1.5 × 10^5^ spores.mL^-1^ and 2.0 × 10^5^ spores.mL^-1^, for which the correlation was strong (r > 0.65). We investigated whether the proportion of virulent strains in the tested *Z. tritici* population could be deduced from ρ_L,_ assuming that the relationship between the two variables is linear with a zero intercept (Figure 7). We obtained *a* = 86.70 with r^2^ = 0.630 for counts on the 5-cm leaf sections and *a* = 97.16 with r² = 0.782 for counts on whole leaves.

**Figure 7.**
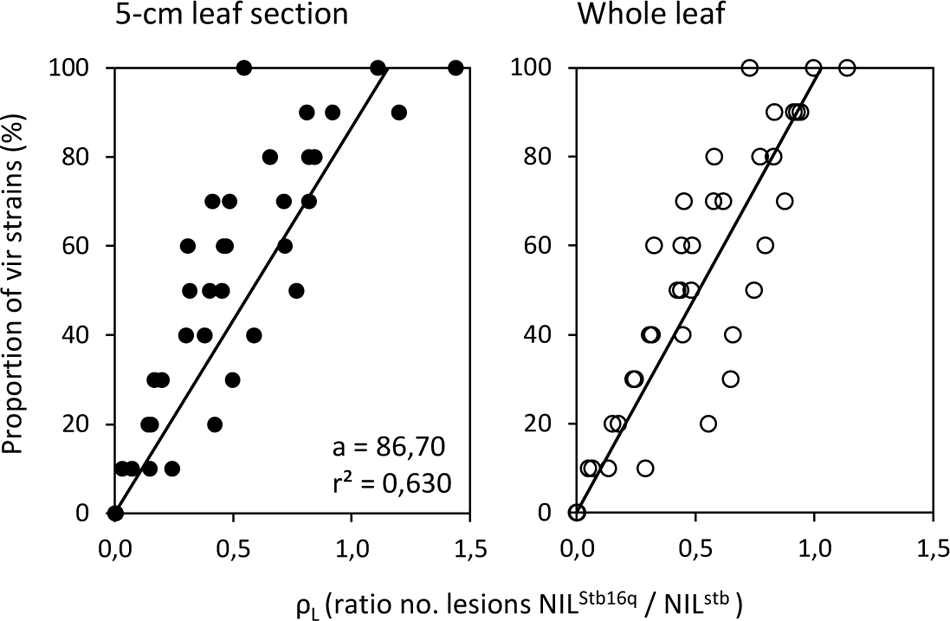
Standard curves for the proportion of *Zymoseptoria tritici* vir strains as a linear function of the ratio (ρ_L_) of the mean number of STB lesions counted on 5-cm leaf section (black points) or whole leaves (white points) of wheat lines NIL^Stb16q^ and NIL^stb^ in a microcanopy obtained with inoculum concentrations of 1.5 × 10^5^ spores.mL^-1^ and 2.0 × 10^5^ spores.mL^-1^ (pooled data set for Mix1A and Mix1B, binary vir-avr mixtures; Table 1). Each line corresponds to the linear regression with a zero intercept, *y = a.x*, where *a* is the estimated slope and r^2^ is the coefficient of determination.

In the second experiment we established ‘standard curves’ for low proportions (below 0.2) of virulent strains with a synthetic population consisting of 10 vir and 10 avr *Z. tritici* strains (Mix2; Table 1). Using six proportions of vir strains (0, 0.05, 0.1, 0.2 and 0.5) for the linear regression analysis, we showed that the slope estimate (*a*) was very close to 100 (range: 93 to 99), whether we used counts of STB lesions on 5-cm leaf sections or on whole leaves (Figure 8B). This result demonstrates that it is feasible to determine the proportion of virulent strains in a diverse population of strains with different levels of aggressiveness with this method.

**Figure 8.**
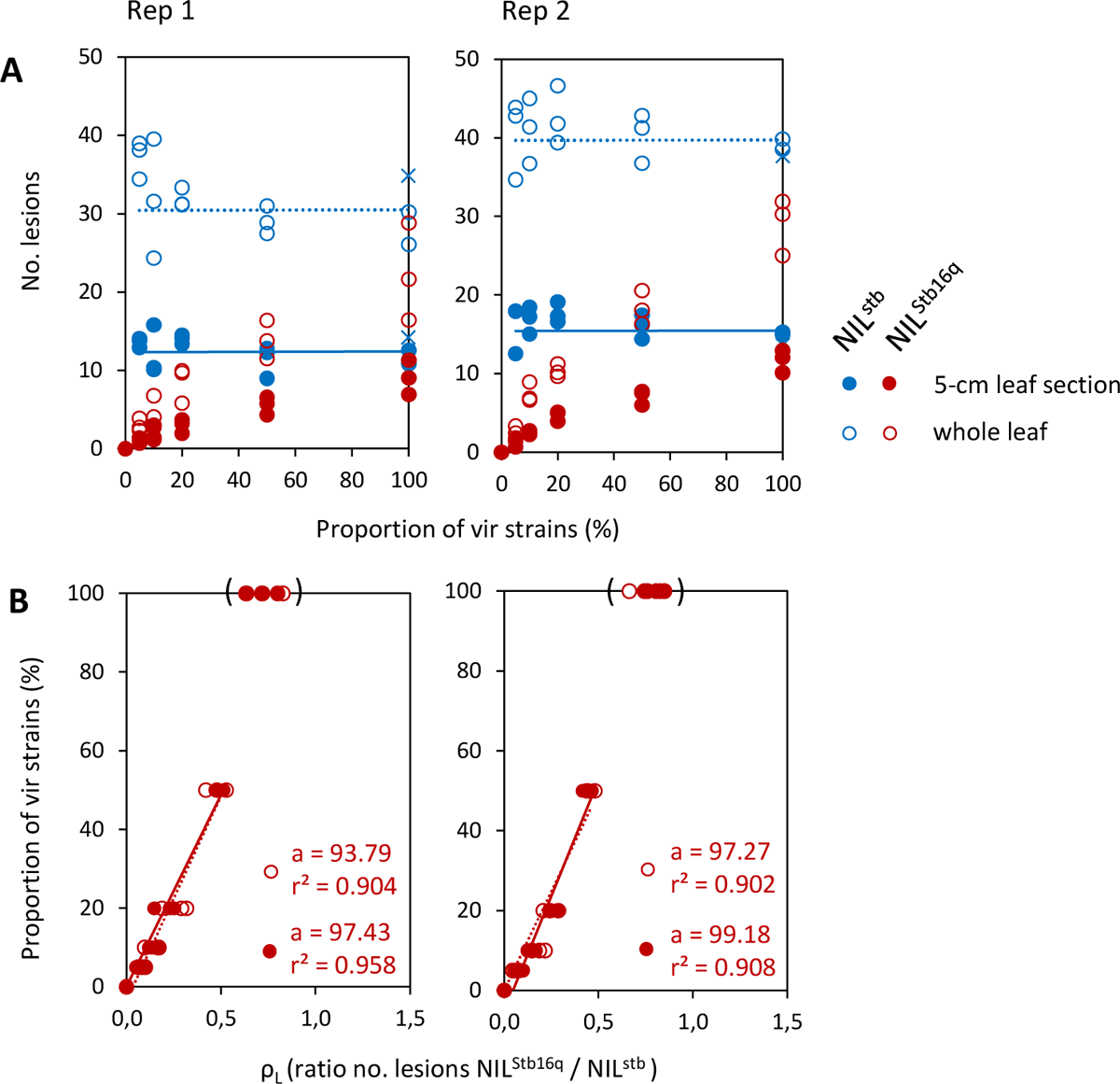
A. Mean numbers of STB lesions counted on 5-cm leaf sections and on whole leaves of wheat lines NIL^stb^ (blue) and NIL^Stb16^ (red) in a microcanopy according to the proportion of *Zymoseptoria tritici* vir strains in the inoculum suspension (Mix2, synthetic population consisting of 10 vir and 10 avr strains; Table 1), for the first (rep 1) and second (rep 2) replicates. Each point is the mean number of lesions on 24 marked leaves. Horizontal blue lines represent the mean number of lesions that appeared on NIL^stb^ whatever the percentage of vir strains in the inoculum suspension. **B.** Standard curves indicating the proportion of vir strains as a linear function of the ratio (ρ_L_) of the mean number of STB lesions on 5-cm leaf sections (black points) or whole leaves (white points) for wheat lines NIL^Stb16^ relative to NIL^stb^ in a microcanopy for rep 1 and rep 2. Each line corresponds to the linear regression with a zero intercept, *y = a.x*, where *a* is the estimated slope and r^2^ is the coefficient of determination (data set from inoculum suspensions containing only vir strains not taken into account).

### Application to a diversified population with an expected composition

In the third experiment, which involved the progenies of the cross between strains FS5657 (vir) and FS2548 (avr), ρ_L_ was close to 0.5 (0.47 in rep 1 and 0.45 in rep 2; Table 2), slightly lower than the ρ_L_ values obtained for the control replicates (0.54 in rep1 and 0.50 in rep 2). This showed 1:1 segregation for vir and avr, consistent with the expected value, further validating the method.

**Table 2.**
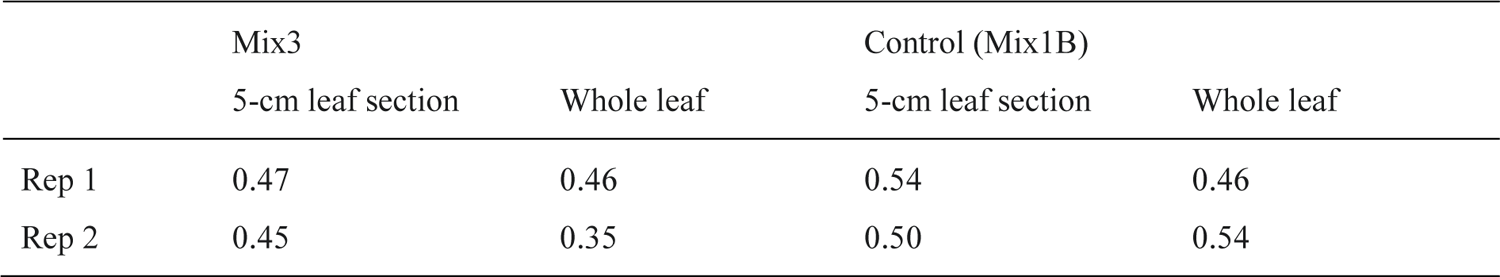
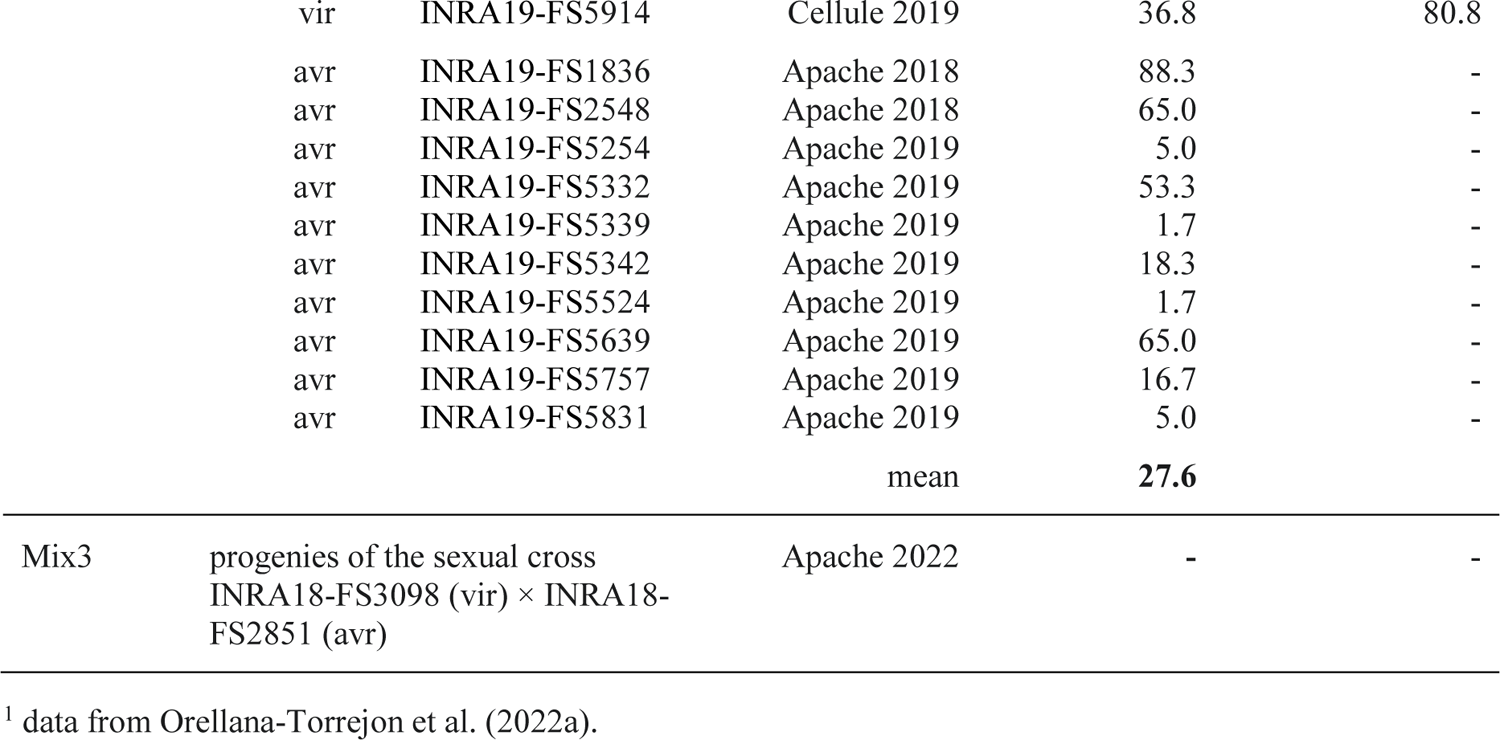
Frequency of virulent strains in the progeny of the sexual cross between strains FS3098 (vir) and FS2851 (avr) (Mix3) estimated by calculating the ratio (ρ_L_) of the number of STB lesions counted on NIL^Stb16q^ to that on NIL^stb^, with Mix1B (FS5657 and FS2548) as the control.

## Discussion

In this study, we validated the principle underlying the ‘bulk phenotyping on a checkerboard microcanopy’ (BPC) method by demonstrating the existence of a strong correlation between the ratio (ρ_L_) of the mean number of STB lesions on a wheat line with the resistance gene to that on a wheat line without the resistance gene and the proportion of virulent strains in the inoculum suspension. ρ_L_ is a more reliable indicator of the proportion of virulent strains in a bulk when the coefficient of determination is high and the relationship is close to proportionality (r^2^ > 0.90 and *a* > 0.97 for 5-cm leaf sections; Figure 8). Our protocol cannot estimate the mean number of spores actually deposited on the leaves or the rate of successful leaf penetration (infection efficiency), and we assumed that the status of a strain (vir or avr) has no effect on infection efficiency in the tissues of a wheat line not carrying the corresponding resistance gene. The testing of this hypothesis would require microscopy observations such as those carried out by Battache et al. (2022), but the confirmation of this hypothesis does not appear to be essential for further application of the BPC method. The method is based on standard curves, the slope (*a*) – but not robustness (r^2^) – of which would probably be affected if this hypothesis did not apply, given that all the ρ_L_ ratios would be influenced identically by differential infection between vir and avr strains.

The standard curve can be used as a reference for quantifying the frequency of virulent strains against *Stb16q* in a field population. A greater deviation of the slope (*a*) from 100 (strict proportionality) would have suggested the existence of methodological biases, or even invalidated the very principle of the BPC method. Non-proportionality was apparent in the preliminary standard curves established with the full range of virulence proportions (0-100; Figures 6), probably due to the effect of inoculum concentration, which proved to be a crucial parameter of the BPC method. Preliminary tests have shown that the concentration must be adjusted carefully, within the range of 1.5 × 10^5^ spores.mL^-1^ to 2.0 × 10^5^ spores.mL^-1^, and adapted according to the facilities and operators. The concentration should not be too high, to prevent saturation, in terms of lesion numbers on the leaves of the wheat line not carrying the *Stb* gene, but also because high inoculum concentrations may mask deficiencies in virulence on the wheat line carrying the *Stb* gene (Fones et al., 2015). Conversely, inoculum concentration should not be too low, to ensure that there are sufficient lesions for counting on the leaves of the line carrying the resistance gene treated with suspensions with low proportions of virulent strains. The lower limit for the quantification of virulence proportions depends on the accuracy of spore concentration adjustment; the lowest proportion of virulence strains used here (0.05) appears to be above this limit, but its quantification might have become less reliable below this concentration. Non-proportionality may also result from interactions between strains during infection, as highlighted in studies focusing on mixed infections (Susi et al., 2015; Tollenaere et al., 2016; Bernasconi et al., 2022; 2023). The possibility of infection by ‘stowaway strains’ through facilitation mechanisms, as in the systemic induced susceptibility reaction, has been suggested for *Z. tritici* (Seybold et al., 2020), consistent with the positive effect of population diversity on symptom intensity sometimes observed (Zelikovitch and Eyal, 1991). Recent reports by Barrett et al. (2021) and Bernasconi et al. (2023) have confirmed that strains virulent against *Stb6* can manipulate the plant immune response in mixed infections, facilitating apoplast colonization by avirulent strains. This mechanism could lead to lesions being caused by avirulent strains on the leaves of the wheat lines carrying the resistance gene, resulting in a ρL ratio slightly higher than the proportion of virulent strains (p) in the inoculum suspension. This contrasts with our findings for *Stb16q*, as the slope was marginally below 1, specifically ρ_L_ < p (Figure 7), which does not call the BPC method into question. Nonetheless, it might be useful to delve deeper into the phenotypic expression of these facilitation mechanisms for other *Stb* genes by conducting re-isolation from the lesions on the wheat line carrying the resistance gene (to check for the presence of avirulent strains).

The BPC method can be used directly to estimate the frequency of *Z. tritici* strains virulent against *Stb16q* in a pathogen population without to the need for individual (strain-by-strain) phenotyping, as described by Orellana-Torrejon et al. (2022a, 2022b). The method would benefit for generalization for virulence against other *Stb* genes for which isolines (e.g. NILs derived from Chinese Spring) are available or for commercial varieties carrying different *Stb* genes (i.e. in which the *Stb* genes have been identified). Such wheat varieties differ by one – or more – *Stb* gene, but also have different genetic backgrounds (e.g. cv Cellule vs. Apache in the case of *Stb16q*; Orellana-Torrejon et al., 2022a, 2022b). For this reason, a relationship between the ρ_L_ ratio and the frequency of virulent strains further removed from proportionality might be expected if commercial varieties were used, and it would be necessary to establish specific standard curves and test their robustness.

The BPC method could be used to monitor the frequency of several virulences at wide spatiotemporal scales or in analytical field experiments. Such monitoring is increasingly relying on molecular methods, and some such methods are currently being developed for STB (e.g. Samils et al. 2021; Gutierrez Vazquez et al. 2022; Bellah et al. 2023a). These methods, based on deep-sequencing technologies with short reads (i.e. Illumina) or long reads (i.e. PacBio, Nanopore), are powerful tools. They enable rapid, cost-effective analyses of large numbers of samples, for a large number of target genes. Sequencing-based methods also have the advantage of revealing allelic diversity within the populations analyzed. However, these methods are constrained by the need to identify the (a)virulence gene associated with the targeted resistance and to link observed allelic polymorphisms in the (a)virulence gene with the avirulent or virulent phenotype. The BPC method is valuable and potentially complementary to sequencing-based approaches, as it does not require prior knowledge of the (a)virulence or resistance gene. In addition, the phenotype is inferred directly from observations.

The BPC method could theoretically be adapted to a broad range of plant pathosystems, but its application is more readily feasible in cases in which the host-pathogen interaction is qualitative. This is the case for *Stb6* (Zhong et al., 2017; Saintenac et al., 2018), *Stb9* (Amezrou et al., 2023) and *Stb16q* (Saintenac et al., 2021), but is the exception rather than the rule in the wheat-*Z. tritici* pathosystem. Indeed, most interactions in this pathosystem are quantitative in nature (Langlands-Perry et al., 2023; Meile et al., 2017; 2023). The validity of the BPC method for predominantly quantitative interactions remain to be determined.

The principle of the BPC method could also theoretically be in field conditions with more precise disease scoring than is usual – numbers of lesions rather than estimated diseased leaf areas – to obtain the frequency of virulent strains in a local population directly, by calculating the difference or the ratio (ρ_L_) of the number of lesions between two carefully selected wheat lines grown at the same site. The efforts required for lesion counts, even if methods for increasing the efficiency of such counts have been developed (e.g. Bousset et al., 2016), may also explain why this principle has been little used for direct monitoring in field conditions. The BPC method could be used to investigate the impact of different practices – the use of cultivar mixtures, for example – on the virulence composition of a *Z. tritici* population without the need for strain-by-strain phenotyping *ex situ* as described by Orellana-Torrejon et al. (2022a, b). In the study by Orellana-Torrejon et al., in which STB severity was assessed by eye as the percentage of the leaf area covered by pycnidia, the cultivar Apache (without *Stb16q*) was, on average, 3.5 times more diseased than the cultivar Cellule (carrying *Stb16q*) in pure stands at the start of the epidemic. Using the ratio of disease severity (expressed as the percentage of the leaf area presenting disease) between the two varieties and, thus, following the principle of the BPC method, we can retrospectively estimate that the frequency of strains virulent against *Stb16q* was locally ±28% (1 / 3.5 × 100). This frequency is twice that estimated (±13 %) by the individual phenotyping of several hundred strains sampled from the field trial. This difference highlights the importance of using an accurate proxy of infection points to calculate the ratio ρ_L_, i.e. a number of lesions rather than a visual assessment of the diseased leaf area. The overestimation may also be partly due to the differences in genetic background between Apache and Cellule, and secondary infections that may already have begun to be expressed at the stage at which the disease was scored. In conclusion, field conditions are a potentially relevant framework in which the principle of the BPC method can be applied, but the response obtained under natural conditions is less precise than that obtained under controlled conditions. The application of this method to field conditions raises practical considerations, including, in particular, the challenge of determining the optimal timing for assessment, given that, in the field, the differences in STB severity resulting from differences between varieties can be amplified by the polycyclic nature of the epidemic. Exploring the application of deep learning tools in plant phenotyping could help overcome certain technical challenges.

## Supporting information

Supplementary_material_full_dataset

## Acknowledgments

We thank Iris Bertiner, Béatrice Beauzone and Guillaume Lucas (INRAE BIOGER, Palaiseau, FR) for technical assistance.

